# Similarity Estimation Between DNA Sequences Based on Local Pattern Histograms of Binary Images

**DOI:** 10.1101/016089

**Authors:** Yusei Kobori, Satoshi Mizuta

**Affiliations:** Graduate School of Science and Technology, Hirosaki University, 3 Bunkyo-cho, Hirosaki, Aomori 036-8561, Japan

**Keywords:** genome sequence, mitochondria, bitmap image, occurrence frequency

## Abstract

Graphical representation of DNA sequences is one of the most popular techniques of alignment-free sequence comparison. In this article, we propose a new method for extracting features of DNA sequences represented by binary images, in which we estimate the similarity between DNA sequences by the frequency histograms of local bitmap patterns on the images. Our method has linear time complexity for the length of DNA sequences, which is practical even for comparison of long sequences. We tested five distance measures to estimate sequence similarities and found that *histogram intersection* and *Manhattan distance* are most appropriate for our method among them.

## 1 Introduction

Sequence alignment^1,2^ is generally used to estimate similarities between relatively short sequences such as nucleotide sequences of genes or amino acid sequences of proteins. However, the time complexity of the alignment is *O*(*L*^2^) for sequences of length *L*, which requires an enormous amount of computation time when *L* is large. Therefore, it is necessary to develop so called *alignment-free* methods, which are independent of alignment, to compare long sequences such as whole genome sequences in practical time.

One of the most popular methods for the alignment-free sequence comparison is *graphical representation* of biological sequences^3^. So far various methods based on the graphical representation have been introduced by many authors. The basic procedure is common to almost all the methods, which is outlined as follows: first, each type of bases in a DNA sequence is replaced by an individual vector on a certain dimensional expression space, two-dimension^4–22^, three-dimension^23–36^, or more^37–40^; next, the vectors are connected successively, drawing a trajectory on the expression space; lastly, the distances between the trajectories, or graphs, are calculated based on a pre-defined distance measure.

In this article, we propose a new method for sequence comparison categorized in the graphical representation. We express a DNA sequence as a binary image—each pixel of a binary image is plotted in either black or white—on a 2-dimensional space and count the occurrence frequencies of 3×3 bitmap patterns on the binary image. The distance between the binary images is measured based on the occurrence frequency histograms of the bitmap patterns. As for distance measures between histograms, we selected five frequently used measures: histogram intersection^41^, Manhattan distance, Bhattacharyya distance^42^, Jensen-Shannon divergence^43^, and Kendall’s rank correlation coefficient^44^. Based on phylogeny of 31 mitochondrial genome sequences, we seek for the most appropriate distance measure for our method among the five.

## 2 Methods

### 2.1 Generating a binary image from a DNA sequence

Here, we describe, step-by-step, the procedure to generate a binary image from a DNA sequence.

#### 2.1.1 Graphical representation of a DNA sequence

At first, we assign two-dimensional vectors that are perpendicular to each other to individual types of bases, A, T, G, and C. The number of the independent variations of the assignment is 3!/2 = 3, when we identify the assignments that can be transformed from each other by the rotations of 90-degree or the inversion with respect to the vertical or horizontal axis (Fig. 1). We chose the left most assignment in Fig. 1, where nucleotides A and T are placed on the upper quadrants, and G and C on the lower ones, so that the GC content of a DNA sequence can be grasped easily from the resultant graphical representation. Next, connecting consecutively the vectors assigned to the bases of the DNA sequence from the origin to the end one by one, we can draw a 2-dimensional graph. Fig. 2(a) shows an example—a graphical representation of sequ

**Figure 1:**
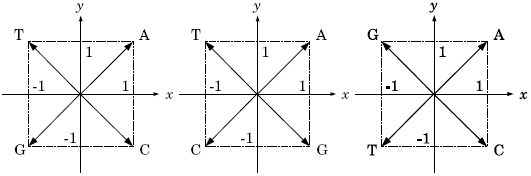
Three independent assignments of vectors to individual bases.

**Figure 2:**
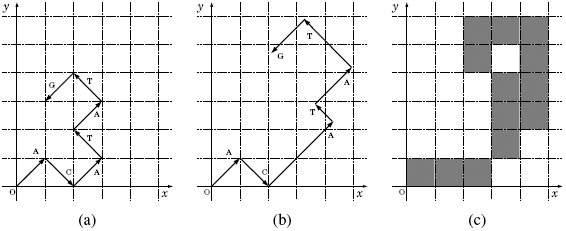
Generating a binary image of sequence “ACATATG”: (a) the primary graphical representation, (b) the graphical representation modified with weighting factors, and (c) the generated binary image.

#### 2.1.2 Multiplying weighting factors

In order to extract potential information conveyed by individual bases, we introduced weighting factors based on a Markov chain model into the process of generating binary images^45^. As the weighting factor, we used self-information *I(E)*, which is the amount of information that we will receive when a certain event *E* occurs. Let *P(E)* be the probability that event *E* occurs, then *I(E)* is defined as *I(E)* = - log_2_ *P(E)* in bits. Here, we define *P(E)* according to the second order Markov chain, concerning about codons in the coding regions of DNA sequences, which is calculated by the occurrence frequencies of triplets of bases. The probability that nucleotide *z* occurs after a pair of nucleotides *xy (x, y,z* ∈ {A, T, G, C}) is calculated by

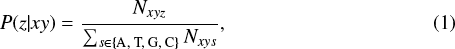

where *N*_*xys*_(*s* ∈ {A, T, G, C}) is the number of occurrence of triplet *xys*, which is measured in all the DNA sequences analyzed.

The individual vectors corresponding to the bases on a DNA sequence are multiplied by the weighting factors according to the preceding two bases. For example, when P(A|AC), P(T|CA), P(A|AT), P(T|TA), and P(G|AT) are .20, .66, .41, .31, and .44, respectively, the weighting factors are calculated to be 2.3, 0.60, 1.3, 1.7, and 1.2, respectively. The first two of a series of vectors in Fig. 2(a) are drawn without weightings because the corresponding weighting factors can not be available. The third vector A is multiplied by 2.3 because the preceding doublet of bases is AC and, therefore, the weighting factor corresponding to P(A|AC) is chosen. The remaining vectors are similarly multiplied by the weighting factors. As a result, the graphical representation of sequence “ACATATG” is modified as shown in Fig. 2(b).

#### 2.1.3 Generating a binary image

A binary image is a digitized image that has pixels of only two possible values 0 and 1, which are typically plotted in *white* and *black*, respectively. From the graphical representation of a DNA sequence, we generate a binary image by the following manner; we set value 1 for the pixels that include at least a part of a vector, and 0 otherwise, in the graphical representation (Fig. 2(c)).

### 2.2 Local patterns

We define a local pattern as a bitmap of a set of adjacent pixels of a certain size of window. Because each pixel of a binary image has two pixel values, the number of local patterns is 2^*n*^, where *n* is the number of pixels belonging to a window. Windows of too large size are dominated by white pixels, and, on the other hand, those of too small size can not have enough variations to express a DNA sequence. In this study, therefore, we chose the window size of 3 × 3, where the number of the local patterns is 2^9^ = 512. Note that we do not include the local pattern whose pixels are all white into the local pattern histograms because it represents the empty background of the images.

Fig. 3 shows the five examples of local patterns of window size 3×3 with their serial numbers below. We derive the serial numbers by lining up the pixels from the upper left corner to the lower right and interpreting them as a binary number with zeros and ones for white and black pixels, respectively, with the upper left corner being the highest bit.

**Figure 3:**
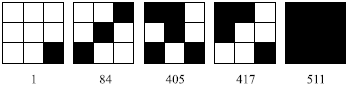
Five examples of local patterns with their serial numbers below. The definition of the serial numbers is described in Subsection 2.2.

### 2.3 Counting the occurrence frequencies of local patterns

Sliding a 3 × 3 window by one pixel per move over the binary image, we counted the occurrence frequencies of the local patterns. If the trajectory of the graphical representation of a DNA sequence is like a *random walk*, the average distance between the origin and the terminus of the trajectory would be *O*(*L*^1/2^), where *L* is the sequence length. In that case, the rectangle area covering the whole trajectory is proportional to *L*; hence the computational time to count the occurrence frequencies would become *O(L)*. In reality, however, the trajectory is not a random walk but a curved line of length *O(L)* in most cases (see Fig. 4 in Section 3), in which the computational time to count the occurrence frequencies becomes *O*(*L*^2^)—this is equivalent to that for pair-wise sequence alignment.

Accordingly we devised the method of counting the occurrence frequencies as follows. We divided the binary image into square regions of 10 × 10 pixel, and marked the regions having at least one black pixel, simultaneously when generating the binary image. Then, we counted the occurrence frequencies of the local patterns only in the marked regions. Thus we can reduce the computational time of counting to *O(L)*, although there remains room for improvement in reducing the numerical coefficients.

### 2.4 Distance measures between local pattern histograms

There are several measures to estimate similarity/dissimilarity between two histograms. Among them, we chose commonly used five measures as the distance measure between the local pattern histograms, and compared them, seeking for the appropriate measure for our method. We briefly describe them below. In the following formulas, *p*_*i*_ and *q*_*i*_ are the occurrence frequencies of the local pattern of serial number _i_ in histograms *P* and *Q*, respectively, and *N* is the largest serial number of the local patterns (i.e., *N* = 511 for 3×3 pixel local patterns). Note that, in the calculation of the distances, the occurrence frequencies are normalized to be 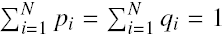.

**Histogram intersection (HI)** Histogram intersection was proposed by Swain *et al*.^41^ for color indexing of images, which is defined as

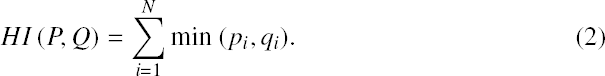

It ranges from 0 to 1, with 1 for *P* and *Q* being identical. To calculate distances, we convert it to *D*_HI_ *(P, Q)* = 1 – *HI (P, Q)*.

**Manhattan distance (MD)** Manhattan distance, also known as *City block distance* or *L*_1_ -*norm*, is defined as

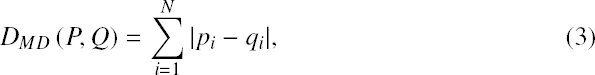

which ranges from 0 to 2, with 0 for *P* and *Q* being identical.

**Bhattacharyya distance (BD)** Bhattacharyya distance ^42^ is defined between two probability distributions from a divergence

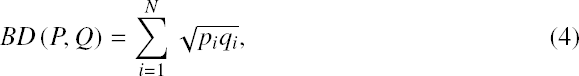

which ranges from 0 to 1, with 1 for *P* and *Q* being identical. The Bhattacharyya distance is defined from the divergence as *DBD (P, Q)* = – ln *BD (P, Q)*.

**Jensen-Shannon divergence (JS)** Jensen-Shannon divergence^43^ is a symmetrized and smoothed version of Kullback-Leibler divergence^46^, which is defined as

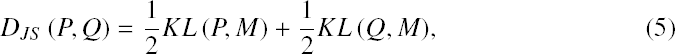

where *M* = (*P + Q*)/2 and *KL* ( ·, *M*) is the Kullback-Leibler divergence calculated by

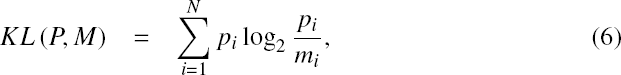

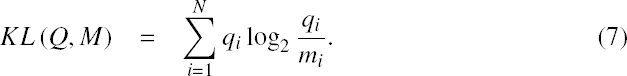

Here, *m*_*i*_ = *(p_i_ + q_i_)*/2. The Jensen-Shannon divergence ranges from 0 to 1, with 0 for *P* and *Q* being identical.

**Kendall’s rank correlation coefficient** (τ) Kendall’s rank correlation coefficient^44^, also known as Kendall’s τ, is defined as

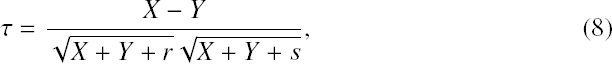

where *X* is the number of *concordant i, j (i > j)* pairs in which *(p_i_ - p_j_)(q_i_ - q_j_)* > 0 is satisfied; *Y* is the number of the *discordant* pairs in which *(p_i_ – p_j_)(q_i_ – q_j_)* < 0 is satisfied; *r* is the number of one kind of the *tie* pairs in which *p*_*i*_ = *p*_*j*_ and *q*_*i*_ ≠ *q*_*j*_ are satisfied; and *s* is the number of the other kind of the *tie* pairs in which *p_i_ ≠ p_j_* and *q*_*i*_ = *q*_*j*_ are satisfied. If both *p*_*i*_ = *p*_*j*_ and *q*_*i*_ = *q*_*j*_ are satisfied, the corresponding *i, j* pairs are excluded from the computation. Kendall’s τ lies between −1 and 1, with 1 for the rank orders of *p*_*i*_*s* and *q*_*i*_*s* being completely in agreement with each other, and with −1 for them being completely reversal with each other. We re-scaled the Kendall’s τ as

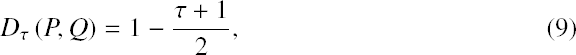

so that *D*_*τ*_ (*P, Q*) ranges from 0 to 1, with 0 for the rank orders of *P* and *Q* being identical.

## 3 Results and Discussion

### 3.1 Genome sequences analyzed

We downloaded mitochondrial genome sequences of 31 mammalian species from GenBank^47^ and analyzed them. Table 1 summarizes the genome sequences. Mitochondrial genomes are widely used to study genome evolution and phylogenetic inference due to, for example, a high mutation rate relative to nuclear genomes and a nearly uniform size for mammalian species.

**Table 1:** Mitochondrial genomes analyzed.

| Ac. No. | Species | Length (bp) |
| --- | --- | --- |
| V00662 | Human | 16569 |
| D38116 | Pygmy chimpanzee | 16563 |
| D38113 | Common chimpanzee | 16554 |
| D38114 | Gorilla | 16364 |
| X99256 | Gibbon | 16472 |
| Y18001 | Baboon | 16521 |
| AY863426 | African green monkey | 16389 |
| D38115 | Bornean orangutan | 16389 |
| U20753 | Cat | 17009 |
| EF551003 | Tiger | 16990 |
| EF551002 | Leopard | 16964 |
| U96639 | Dog | 16727 |
| EU442884 | Wolf | 16774 |
| AJ002189 | Pig | 16680 |
| AF010406 | Sheep | 16616 |
| V00654 | Cow | 16338 |
| AY488491 | Buffalo | 16355 |
| X97336 | Indian rhinoceros | 16829 |
| Y07726 | White rhinoceros | 16832 |
| X63726 | Harbor seal | 16826 |
| X72004 | Gray seal | 16797 |
| AJ224821 | African elephant | 16866 |
| DQ316068 | Asiatic elephant | 16902 |
| DQ402478 | Black bear | 16868 |
| AF303110 | Brown bear | 17020 |
| AF303111 | Polar bear | 17017 |
| AJ001588 | Rabbit | 17245 |
| X88898 | Hedgehog | 17447 |
| X14848 | Norway rat | 16300 |
| AF348082 | Vole | 16312 |
| AJ238588 | Squirrel | 16507 |

### 3.2 Weighting factors

We counted the number of occurrences of every tri-nucleotides in all the genome sequences listed in Table 1, sliding a window of length three by one nucleotide per move, and calculated the weighting factors by the method described in Section 2.1. The calculated weighting factors are shown in Table 2. The large (small) weighting factors indicate that the corresponding triplets rarely (often) occur in the genome sequences.

**Table 2:** Calculated weighting factors.

| 1st base | 2nd base |  |  |  | 3rd base |
| --- | --- | --- | --- | --- | --- |
|  | A | C | G | T |  |
| A | 1.64 | 1.66 | 2.08 | 1.68 | A |
|  | 1.90 | 1.80 | 1.58 | 1.94 | C |
|  | 2.94 | 3.28 | 2.22 | 2.85 | G |
|  | 1.84 | 1.76 | 2.23 | 1.79 | T |
| C | 1.68 | 1.80 | 1.82 | 1.37 | A |
|  | 1.91 | 1.70 | 1.83 | 2.03 | C |
|  | 2.88 | 3.57 | 2.43 | 3.01 | G |
|  | 1.80 | 1.65 | 2.00 | 2.03 | T |
| G | 1.57 | 1.70 | 1.57 | 1.33 | A |
|  | 2.07 | 1.51 | 1.87 | 2.23 | C |
|  | 2.30 | 3.95 | 2.45 | 2.73 | G |
|  | 2.17 | 1.86 | 2.27 | 2.07 | T |
| T | 1.71 | 1.54 | 1.46 | 1.55 | A |
|  | 2.01 | 1.82 | 2.12 | 1.87 | C |
|  | 2.56 | 3.38 | 2.31 | 3.02 | G |
|  | 1.85 | 1.85 | 2.29 | 1.94 | T |

### 3.3 Graphical representations

Fig. 4 shows the graphical representations of the 31 mammals. It can be recognized that the trajectories among closely related species, such as primates, cats, elephants, bears, and so on, look similar to each other; on the other hand, those between the species in different orders look different. This observation indicates that our method of graphical representation is effective in visual inspection of the sequence similarities. The usefulness of introducing the weighting factors in the graphical representation is argued by one of the authors^45,48^.

**Figure 4:**
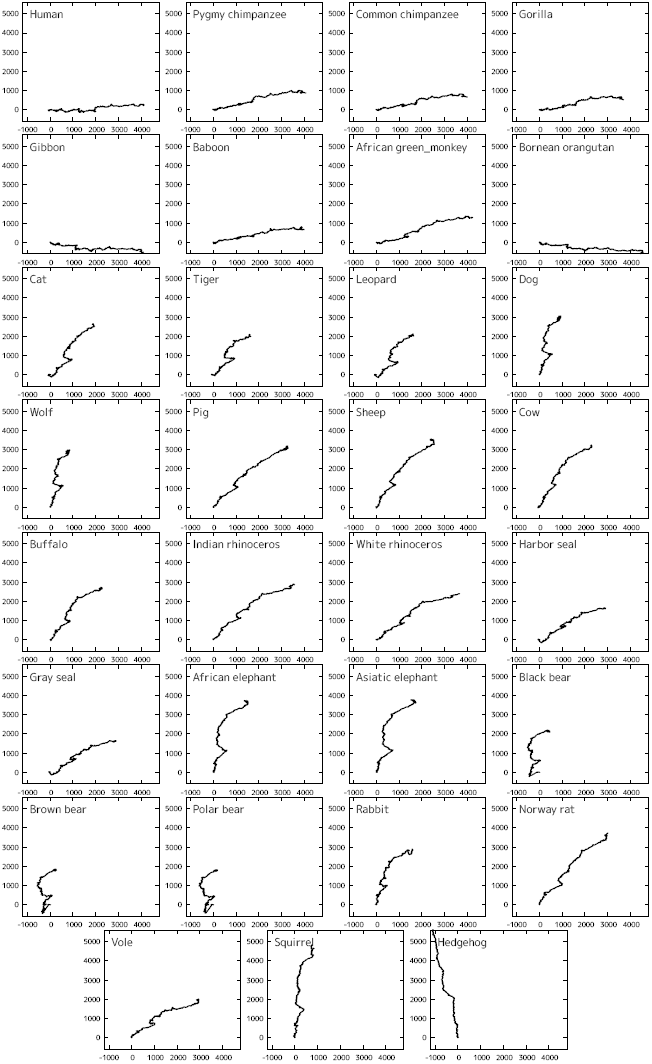
Graphical representation of mitochondrial genomes of 31 mammals.

### 3.4 Local pattern histograms

We counted the numbers of occurrences of the local patterns for the 31 mammalian species and constructed the local pattern histograms. Fig. 5 shows the (un-normalized) local pattern histograms of the 31 mammals. Local patterns 1, 4, 10, 34, 64, 84, 136, 160, 256, and 273 are detected more than 1000 counts in all the genome sequences. Those frequent local patterns are depicted in Fig. 6.

**Figure 5:**
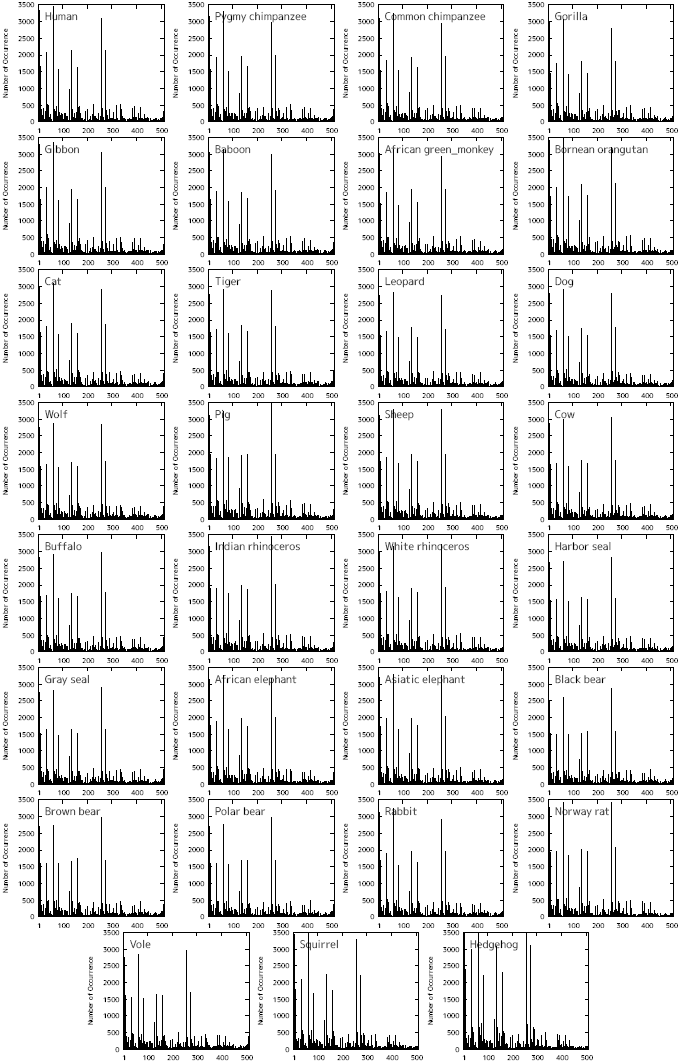
Local pattern histograms of mitochondrial genomes of 31 mammalian species.

**Figure 6:**
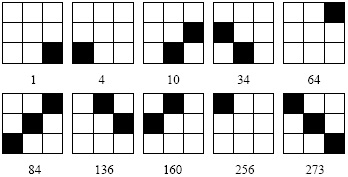
Frequently occurring local patterns with their serial numbers below. The definition of the serial numbers is described in Subsection 2.2.

### 3.5 Construction of phylogenetic trees

Fig. 7 shows the phylogenetic trees constructed from the calculated distance matrices using Unweighted Pair Group Method with Arithmetic mean (UPGMA). The trees are drawn by statistical analysis software R^49^. The trees of the top panel, the middle one, and the bottom one are constructed based on distance measures HI and MD, BD and JS, and τ, respectively. The upper two trees seem to be well reconstructed in that primates, elephants, cats, bears, and so on, are located in their respective clades, although sheep is separated from buffalo-cow pair in the middle tree. In the bottom tree, on the other hand, some species are located on inadequate places; for instance, pig and white rhinoceros are included in primates, and leopard is separated from cat-tiger pair.

**Figure 7:**
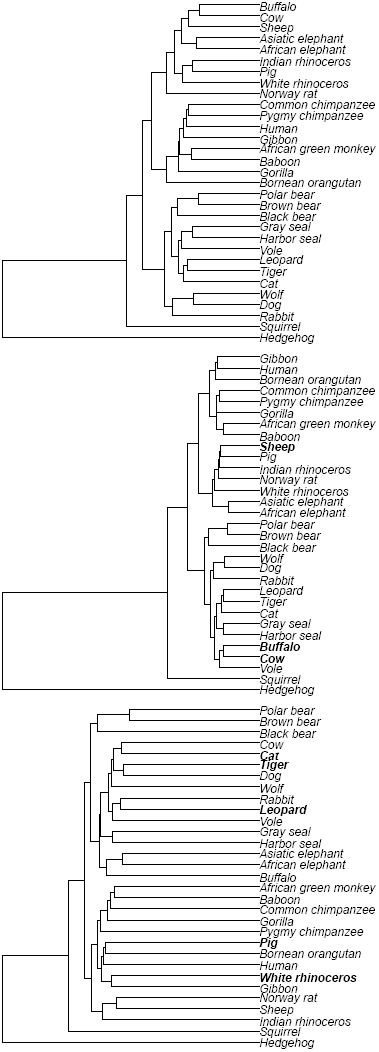
Phylogenetic trees constructed using UPGMA method based on distance measures, histogram intersection and Manhattan distance (top panel), Bhattacharyya distance and Jensen-Shannon divergence (middle panel), and Kendall’s τ (bottom panel).

Table 3 shows the Pearson’s correlation coefficients calculated from the distance matrices among the five distance measures. We can find that HI-MD pair and BD-JS pair are strongly correlated with each other, which confirms the topological aspects of the resultant phylogenetic trees in Fig. 7.

**Table 3:** Pearson’s correlation coefficients among the five distance measures.

|  | HI<sup>a</sup> | MD<sup>b</sup> | BD<sup>c</sup> | JS<sup>d</sup> |
| --- | --- | --- | --- | --- |
| MD | 0.99933 |  |  |  |
| BD | 0.97994 | 0.97277 |  |  |
| JS | 0.98091 | 0.97389 | 0.99997 |  |
| τ<sup>e</sup> | 0.94098 | 0.93999 | 0.93992 | 0.94050 |

<sup>a</sup>Histogram intersection
<sup>b</sup>Manhattan distance
<sup>c</sup>Bhattacharyya distance
<sup>d</sup>Jensen-Shannon divergence
<sup>e</sup>Kendall’s $\tau$

To evaluate our phylogenetic trees quantitatively, we measured the Robinson-Foulds (R-F) distances^50^ between our trees and a reference tree (Fig. 8), which was constructed by ClustalW^51^ based on the multiple sequence alignment of the mitochondrial genome sequences. The R-F distances were calculated by *treedist* program in Phylogeny Inference Package (PHYLIP)^52^. Table 4 summarizes the measured R-F distances. Among the five distance measures, HI and MD show the best performance.

**Figure 8:**
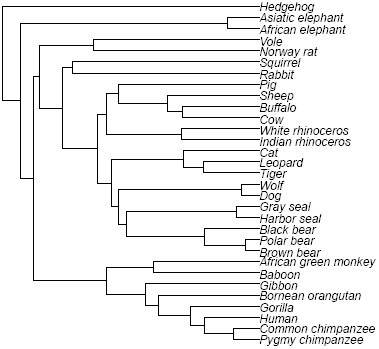
Reference tree constructed by ClustalW.

**Table 4:** Robinson-Foulds distances between phylogenetic trees for the five distance measures.

| Distance measure | R-F distance |
| --- | --- |
| HI<sup>a</sup>, MD<sup>b</sup> | 30 |
| BD<sup>c</sup>, JS<sup>d</sup> | 34 |
| τ<sup>e</sup> | 46 |

<sup>a</sup>Histogram intersection
<sup>b</sup>Manhattan distance
<sup>c</sup>Bhattacharyya distance
<sup>d</sup>Jensen-Shannon divergence
<sup>e</sup>Kendall’s $\tau$

At this point, it is worth discussing the position of hedgehog. We compared our tree constructed by HI and MD (the top panel of Fig. 7) with those given by Huang *et al*.^21^ and Yu *et al*.^53^ Overall configuration of our tree approximately agrees with their trees except for the position of hedgehog. Krettek *et al*.^54^ performed a phylogenetic analysis using concatenated sequences of 13 protein-coding genes of mitochondrial genomes of nine mammals including human, harbor seal, cow, and hedgehog, and identified the position of hedgehog as basal relative to the other species included. Our result is consistent with Krettek *et al*.^54^ rather than Huang *et al*.^21^ and Yu *et al*.^53^

## 4 Conclusion

In this study, we proposed a novel method for estimating similarities between DNA sequences. In this method, we express DNA sequences as binary images by replacing individual bases with 2-dimensional vectors and connecting them successively. We counted the occurrence frequencies of 3 × 3 bitmap patterns on the binary images and measured the distances between them based on the frequency histograms of the bitmap patterns. We tried five frequently used distance measures to estimate similarity/dissimilarity between two histograms: histogram intersection, Manhattan distance, Bhattacharyya distance, Jensen-Shannon divergence, and Kendall’s rank correlation coefficient.

We compared our phylogenetic trees with that constructed by ClustalW for mitochondrial genomes of 31 mammalian species. Among the five distance measures, histogram intersection and Manhattan distance showed the best performance in terms of Robinson-Foulds distance between the phylogenetic trees, as well as having the more appropriate position of hedgehog on the phylogenetic tree than those given by Huang *et al*.^21^ and Yu *et al*.^53^.

The most time consuming step in our method is counting the occurrence frequencies of local patterns. Its time complexity is *O(L)* for sequence length *L*, which is practical even for very long sequences, although the proportionality constant is large; it is thought to be possible to make the constant smaller to reduce the actual computation time by improving the counting method.

